# Sera from individuals with narrowly focused influenza virus antibodies rapidly select viral escape mutations *in ovo*

**DOI:** 10.1101/324707

**Authors:** Amy K.F. Davis, Kevin McCormick, Megan E. Gumina, Joshua G. Petrie, Emily T. Martin, Katherine S. Xue, Jesse D. Bloom, Arnold S. Monto, Frederic D. Bushman, Scott E. Hensley

## Abstract

Influenza viruses use distinct antibody escape mechanisms depending on the overall complexity of the antibody response that is encountered. When grown in the presence of a hemagglutinin (HA) monoclonal antibody, influenza viruses typically acquire a single HA mutation that reduces the binding of that specific monoclonal antibody. In contrast, when confronted with mixtures of HA monoclonal antibodies or polyclonal sera that have antibodies that bind several HA epitopes, influenza viruses acquire mutations that increase HA binding to host cells. Recent data from our laboratory and others suggest that some humans possess antibodies that are narrowly focused on HA epitopes that were present in influenza virus strains that they were likely exposed to in childhood. Here, we completed a series of experiments to determine if humans with narrowly focused HA antibody responses are able to select for influenza virus antigenic escape variants *in ovo.* We identified three human donors that possessed HA antibody responses that were heavily focused on a single HA antigenic site. Remarkably, sera from all three of these donors selected single HA escape mutations during *in ovo* passage experiments, similar to what has been previously reported for single monoclonal antibodies. These single HA mutations directly reduced binding of serum antibodies used for selection. We propose that new antigenic variants of influenza viruses might originate in individuals that produce antibodies that are narrowly focused on HA epitopes that were present in viral strains that they encountered in childhood.

**Importance:** Influenza vaccine strains must be updated frequently since circulating viral strains continuously change in antigenically important epitopes. Our previous studies have demonstrated that some individuals possess antibody responses that are narrowly focused on epitopes that were present in viral strains that they encountered during childhood. Here, we show that influenza viruses rapidly escape this type of polyclonal antibody response when grown *in ovo* by acquiring single mutations that directly prevent antibody binding. These studies improve our understanding of how influenza viruses evolve when confronted with narrowly focused polyclonal human antibodies.

## Introduction

Influenza viruses continuously acquire mutations in antigenically important regions of the hemagglutinin (HA) and neuraminidase (NA) proteins through a process termed ‘antigenic drift’ (1). Most humans are infected with influenza viruses in childhood (2) and then continuously re-infected with antigenically distinct strains later in life (3). Early childhood influenza infections can leave lifelong immunological ‘imprints’ that can influence how an individual subsequently responds to antigenically distinct influenza strains (4, 5). Antibody responses in some individuals can become heavily focused on HA epitopes that are conserved between contemporary viral strains and strains that they encountered in childhood (4). It is thought that memory B cells established from childhood infections are continuously recalled later in life upon infection with new influenza virus strains that possess some shared epitopes (4).

Seasonal influenza vaccines need to be continually updated because of antigenic drift (6). Understanding the specificity of common human anti-influenza virus antibodies is important to guide the selection of appropriate seasonal vaccine strains. Selection of appropriate vaccine strains also requires a deep mechanistic understanding of how influenza viruses escape human antibodies. Multiple studies suggest that influenza viruses utilize distinct escape mechanisms depending on the complexity of the antibody response they encounter (as reviewed in (1)). For example, influenza viruses grown *in vitro* or *in ovo* in the presence of a single HA monoclonal antibody rapidly acquire single HA mutations that prevent the binding of the selecting monoclonal antibody (7). In contrast, viruses grown *in vitro* in the presence of multiple HA monoclonal antibodies targeting distinct epitopes (8) or in the presence of polyclonal sera *in vivo* in mice (9) acquire single HA ‘adsorptive’ mutations that increase viral attachment to host receptors. Adsorptive mutations do not always directly prevent antibody binding—instead, viruses with stronger binding avidity can circumvent antibodies of various specificities by simply binding to cells more efficiently (8, 9).

It is unknown if some humans possess polyclonal antibodies that are so biased towards a single HA epitope that they select for single antigenic escape variants similar to what has been previously reported for single monoclonal antibodies. Here, we used hemagglutination-inhibition (HAI) assays and enzyme-linked immunosorbent assays (ELISAs) to identify three human serum samples that possessed anti-H1N1 HA antibodies that were narrowly focused on an epitope that was conserved in an H1N1 strain that circulated during their childhood. We sequentially passaged the A/California/07/2009 H1N1 strain in the presence of these serum samples *in ovo* and characterized the passaged viruses.

## Materials and methods

### Human Samples

Serum samples were previously collected from human participants in the Household Influenza Vaccine Evaluation (HIVE) cohort that was approved by the University of Michigan Medical School institutional review board (10). The University of Pennsylvania institutional review board approved experiments using de-identified serum samples collected at the University of Michigan.

### Viruses

The HA and NA genes of the A/California/07/2009 H1N1 strain were cloned into the vector pHW2000. The A/California/07/2009 HA used in these studies was engineered to possess the D225G mutation that allows the virus to replicate efficiently in eggs without additional egg-adaptive mutations (11). Viruses with A/California/07/2009 HA and NA and A/Puerto Rico/8/1934 internal genes were produced via reverse-genetics by transfecting co-cultures of 293T and MDCK-SIAT1 cells. Transfection supernatants were collected 3 days after transfection and stored at −80 °C. Viruses from transfection supernatants were further amplified in embryonated chicken eggs. We sequenced the HA of our stock viruses and confirmed that HA mutations did not arise during viral propagation in eggs. For some experiments, we used viruses expressing the A/California/07/2009 HA with a K166Q substitution that was introduced using site-directed mutagenesis. For other experiments, we used site-directed mutagenesis and reverse-genetics to create viruses expressing the A/California/07/2009 HA with substitutions that arose during our passaging experiments.

### Viral Passaging in Embryonated Eggs

For the first viral passage, we incubated different dilutions of each human serum sample (1:10, 1:40, 1:160) with 4×10^5^ TCID50 of virus in a total volume of 50ul for 30 minutes at 23°C. These serum/virus mixtures were injected into day 10 embryonated chicken eggs (1 egg per serum/virus mixture) and allantoic fluid from each egg was collected three days later. Viral titers in allantoic fluid were determined by TCID50 using MDCK cells. For the second viral passage, we used virus isolated from the most concentrated amount of serum that did not lead to complete neutralization during the first passage. For example, if we were able to isolate virus from conditions involving 1:160 and 1:40 dilutions of serum but not a 1:10 dilution of serum in the first passage, then we used virus incubated in the first passage in the presence of 1:40 dilution of serum (i.e. the highest amount of serum without complete neutralization). Similar to the first passage, we incubated different dilutions of each human serum sample (1:10, 1:40, 1:160) with 4×10^5^ TCID50 of virus isolated from passage 1 in a total volume of 50ul for 30 minutes at 23°C. These serum/virus mixtures were injected into embryonated chicken eggs (1 egg per serum/virus mixture) and allantoic fluid was collected three days later. Viral titers in allantoic fluid for the second passage were determined by TCID50 using MDCK cells. We carried out the third passage in the same manner as the second passage.

### Sequencing

Bulk RNA was extracted from allantoic fluid samples using the QIAamp Viral RNA Mini Kit (QIAGEN) according to manufacturer’s instructions. Previously described primers were used for full-length amplification of the influenza A genome. Reverse transcription was performed using Superscript III First-Strand Reaction Mix (Thermo Fisher) and an equimolar mix of the 5’-Hoffmann-U12-A4 and 5’-Hoffmann-U12-G4 primers, which bind to the conserved U12 region present on each influenza gene segment (12). 1uL annealing buffer and 1uL of 2 uM primer mix were added to 6uL RNA eluent, then incubated at 65°C for 5 min. 10 uL 2X First-Strand Reaction Mix and 2uL Superscript III/RNaseOUT Enzyme Mix were added on ice for a 20uL total reaction volume, then incubated at 25°C for 10 min, 50°C for 50 min, and 85°C for 5 min. The entire 20uL volume of the reverse-transcription reaction was used as template in a 100uL PCR reaction using KOD HotStart Reaction Mix (EMP Millipore) and a 24-primer cocktail at a total concentration of 600 nM as previously described (13). 35 cycles of PCR amplification were performed with an annealing temperature of 55°C and an extension time of 3 min. An equimolar mix of plasmids containing A/California/07/2009 HA and NA, and A/Puerto Rico/8/1934 PB1, PB2, PA, NP, NS, and M was amplified simultaneously to estimate PCR, library preparation, and sequencing errors. DNA was purified with 1X AMPure beads (Beckman Coulter) and libraries were created with the Illumina sequencing Nextera XT kit. Libraries were sequenced on a MiSeq platform (Illumina) with 250 bp paired-end reads. Library preparation and sequencing was performed in triplicate, starting from independent reverse-transcription reactions.

### Processing of Sequence Data

We removed from downstream analysis samples that did not have at least 10,000 raw sequencing reads. Raw sequencing reads were mapped to the flu genome with bwa, using default parameters. The samtools package was used to sort and index alignment files (.bam) and then to convert them to .pileup format which were used as input for varscan mpileup2cns (--min-coverage 30, --min-var-freq 0.01) to call mutations. A custom R script calculated each mutation’s amino acid position based on its nucleotide position within each segment.

### Quality Filtering

Because sequencing errors are more common towards the end of each viral gene segment, we manually inspected mutations within control samples and determined positional cutoffs for each segment. For the HA segment, we removed mutations outside of nucleotides 1-1,667; for M, 5-1,011; for NP, 5-1,548; for NS1, 5-874; for PA, 5-2,217; for PB1, 14-2,326; for PB2, 5-2,325. We did not filter mutations on NA based on position. We then removed mutations that did not appear in at least 2 technical replicates within each sample. Finally, we removed mutations with an average allele frequency less than 2.5%, which was the highest average frequency seen in any of the CV (amplicon) samples.

### HAI Assays

Serum samples were pretreated with receptor-destroying enzyme (RDE) (Denka Seiken) and HAI titrations were performed in 96-well round bottom plates (BD). Sera were serially diluted twofold and added to four agglutinating doses of virus in a total volume of 100 μL. Turkey erythrocytes (Lampire) were added [12.5 μL of a 2% (vol/vol) solution]. The erythrocytes were gently mixed with sera and virus and agglutination was determined after incubating for 60 minutes at room temperature. HAI titers were expressed as the inverse of the highest dilution that inhibited four agglutinating doses of turkey erythrocytes. Each HAI assay was performed independently on three different dates.

### Virus-Like Particles

Virus-like particles (VLPs) expressing the A/California/07/09 HA with and without amino acid substitutions were produced. We used HAs that possess a Y98F substitution which prevents HA binding to sialic acid (14). It has previously been shown that Y98F has minimal effects on antigenic structure (15). VLPs with HA that has the Y98F substitution can be produced in the absence of NA, which is advantageous for studying anti-HA antibodies in polyclonal sera. For VLP production, we transfected 293T cells (90% confluent in T175 flask) with plasmids expressing HIV gag (36ug), HAT (human airway trypsin-like protease) (0.54ug), and each HA (3.6ug) using polyethylenimine transfection reagent. We isolated culture supernatants and concentrated VLPs using a 20% sucrose cushion. HA amounts in each VLP preparation were determined via ELISAs using the 146-B09 or 065-C05 monoclonal antibodies which bind to conserved regions of the H1 head and stalk, respectively (16).

### ELISAs

ELISA plates were coated overnight at 4°C with VLPs. ELISA plates were blocked with a 3% (wt/vol) BSA solution in PBS for 2 hours. Plates were washed three times with ddH_2_O, and serial dilutions of each serum sample were added to plates in 1% BSA in PBS. After 2 hours of incubation, plates were washed again and a peroxidase-conjugated anti-human secondary antibody (product number 109-036-098; Jackson ImmunoResearch) or a peroxidase-conjugated anti-mouse secondary antibody (MP Biomedicals) diluted in 1% (wt/vol) BSA solution was added. After a one hour incubation, plates were washed, and a 3,3′,5,5′-tetramethylbenzidine (TMB) substrate (product number 50–00-03; Seracare) was added. HCl was added to stop TMB reactions, and absorbances were measured using a plate reader. One-site specific binding curves were fitted in Prism software. After background subtraction (from ELISA plates that did not have VLPs), serum binding to each VLP relative to the vaccine strain was determined using area under the curve. Equivalent coating of each plate was verified using the 065-C05 or 146-B09 monoclonal antibodies (16).

## Results

### Identification of individuals who possess narrowly focused anti-H1N1 antibodies

We have previously shown that some individuals born before 1985 possess anti-H1N1 antibodies that target an HA epitope that is conserved between the A/California/07/2009 pandemic H1N1 strain and seasonal H1N1 strains that circulated prior to 1985 (11, 17). This particular antibody specificity appears to be important, since contemporary H1N1 strains (which are descendants of the 2009 pandemic strain) have acquired several substitutions in the HA epitope targeted by these antibodies over the past 5 years (Fig. 1A-B). For example, H1N1 viruses acquired a K166Q substitution in this epitope during the 2013-2014 season, a S165N substitution in this epitope during the 2015-2016 season, and a S167T substitution in this epitope during the 2016-2017 season (Fig. 1A-B) (H3 numbering used throughout this manuscript). While it is clear that antibodies targeting H1N1 HA epitopes involving residues 165-167 are unable to recognize H1N1 viruses that are currently circulating among humans, it is unknown whether these antibodies are present in sufficient quantities within single individuals to actually select for HA antigenic escape variants with substitutions at residues 165, 166, and 167.

**Fig 1.**
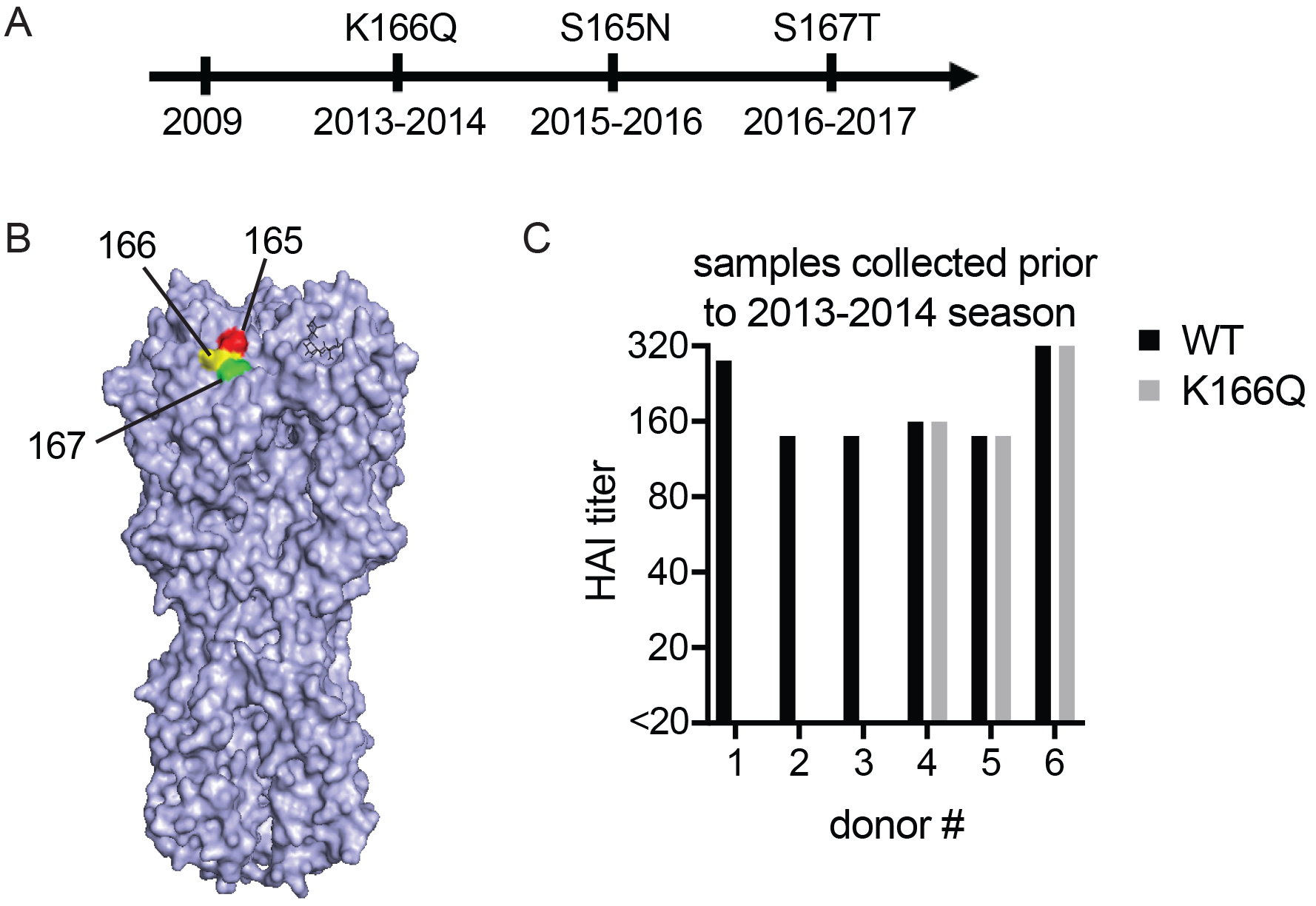
Identification of humans with narrowly focused H1N1 antibody responses. (A) H1N1 viruses acquired substitutions at HA residues 165, 166, and 167 since 2009. (B) Residues 165, 166, and 167 of HA are located in close proximity to each other (PDB ID code 3UBN). (C) HAI assays were completed using H1N1-WT and H1N1-K166Q viruses and serum collected from human donors prior to the 2013-2014 season.

We examined serum samples collected from human donors prior to the 2013-2014 influenza season (10, 17). We identified three donors that were born prior to 1985 who possessed serum antibodies that had >4 fold reduction in hemagglutination-inhibition (HAI) titers using an A/California/07/2009 strain engineered to possess a K166Q HA mutation (herein referred to as H1N1-K166Q) compared to the original ‘wild type’ A/California/07/2009 strain (H1N1-WT) (Fig. 1C). We also identified three donors that possessed serum antibodies that had similar HAI titers using the H1N1-K166Q and H1N1-WT strains (Fig. 1C). HAI assays only detect antibodies that target epitopes near the receptor binding domain in the HA globular head. For that reason, we further characterized each serum sample via ELISAs designed to detect antibodies that bind to both the HA globular head and stalk domains. For ELISA experiments, we engineered a panel of virus-like particles (VLPs) that possess A/California/07/2009 HA with different single point mutations in the HA head and stalk domain (Fig. 2). As expected, serum antibodies from the three donors that reacted poorly with H1N1-K166Q in HAI assays also bound poorly to VLPs possessing HA with the K166Q substitution (Fig. 3A-C). Serum antibodies from donor #1 also had slightly reduced binding to VLPs possessing HA with the N129Y substitution (Fig. 3A) and antibodies from donor #3 had slightly reduced binding to VLPs possessing HA with the K163E mutation (Fig. 3C), both of which are adjacent to HA residue 166 (Fig. 2). Serum antibodies from donors with narrowly focused HAI responses did not have reduced binding to VLPs possessing substitutions in the lower globular HA head and stalk domains (Fig. 3D-F). Serum antibodies from the three donors that had similar HAI titers using the H1N1-K166Q and H1N1-WT viruses did not have reduced binding to any mutant VLPs in our panel, including VLPs possessing HA with the K166Q substitution (Fig. 4A-F). None of the HA substitutions in our VLP panel significantly affected binding of serum antibodies from these three individuals. This could be because these donors possess antibodies that target HA epitopes that are not mutated in our VLP panel. Alternatively, it is possible that these donors have more balanced antibody responses and that our assays are not sensitive enough to detect small reductions in polyclonal antibody binding to each VLP.

**Fig 2.**
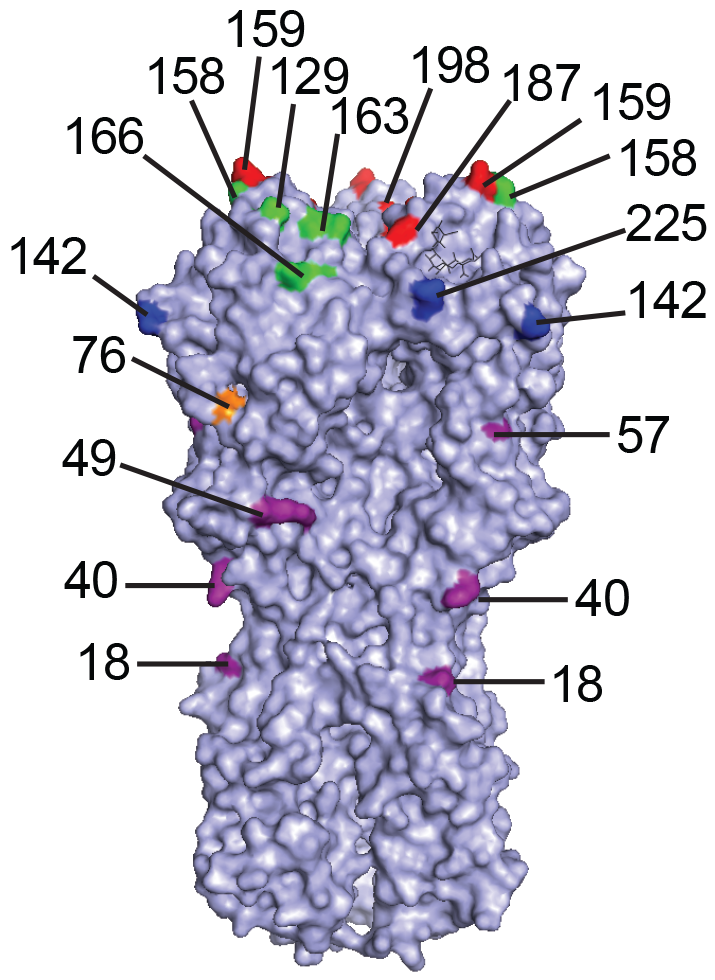
Distribution of substitutions across different HA antigenic sites in our VLP panel. The HA structure (PBD ID code 3UBN) highlighting substitutions in our VLP panel is shown. Sa residues are shown in green, Sb residues are shown in red, Ca residues are shown in dark blue, Cb residues are shown in orange, and lower HA/stalk residues are shown in purple.

**Fig 3.**
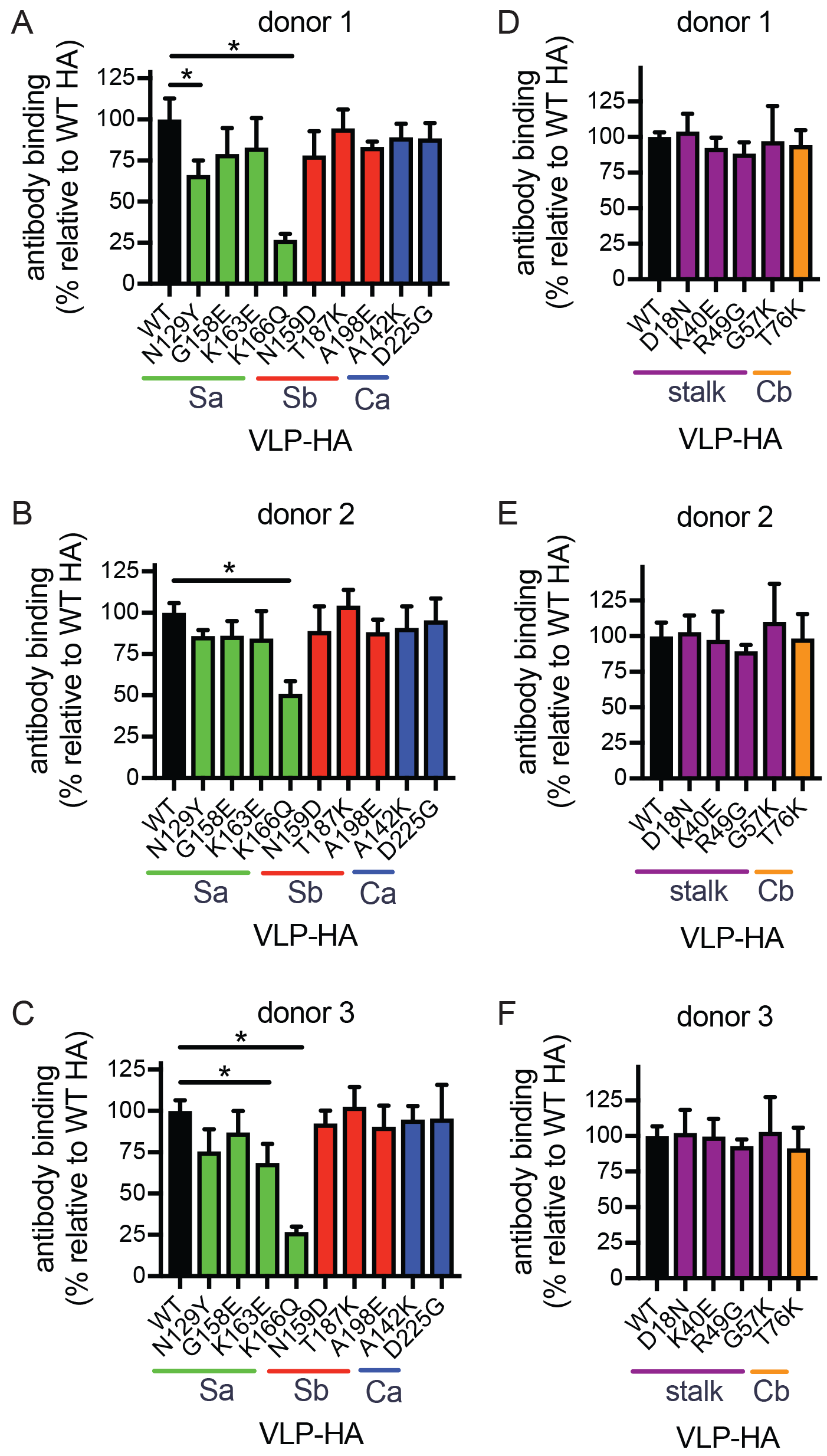
Antigenic characterization of human serum samples that are narrowly focused on a single HA antigenic site. ELISAs were completed using VLPs with HAs that had substitutions in either the top of the HA globular head (A-C) or lower part of HA (D-F). Antibody binding between different ELISA plates was normalized by including a standard monoclonal on each plate. A series of 8 serum dilutions were tested and data are expressed as % binding (based on area under the curve) relative to binding to WT HA. Experiments were completed in triplicate. Mean and SD from three different experiments are shown. (*P<0.05)

**Fig 4.**
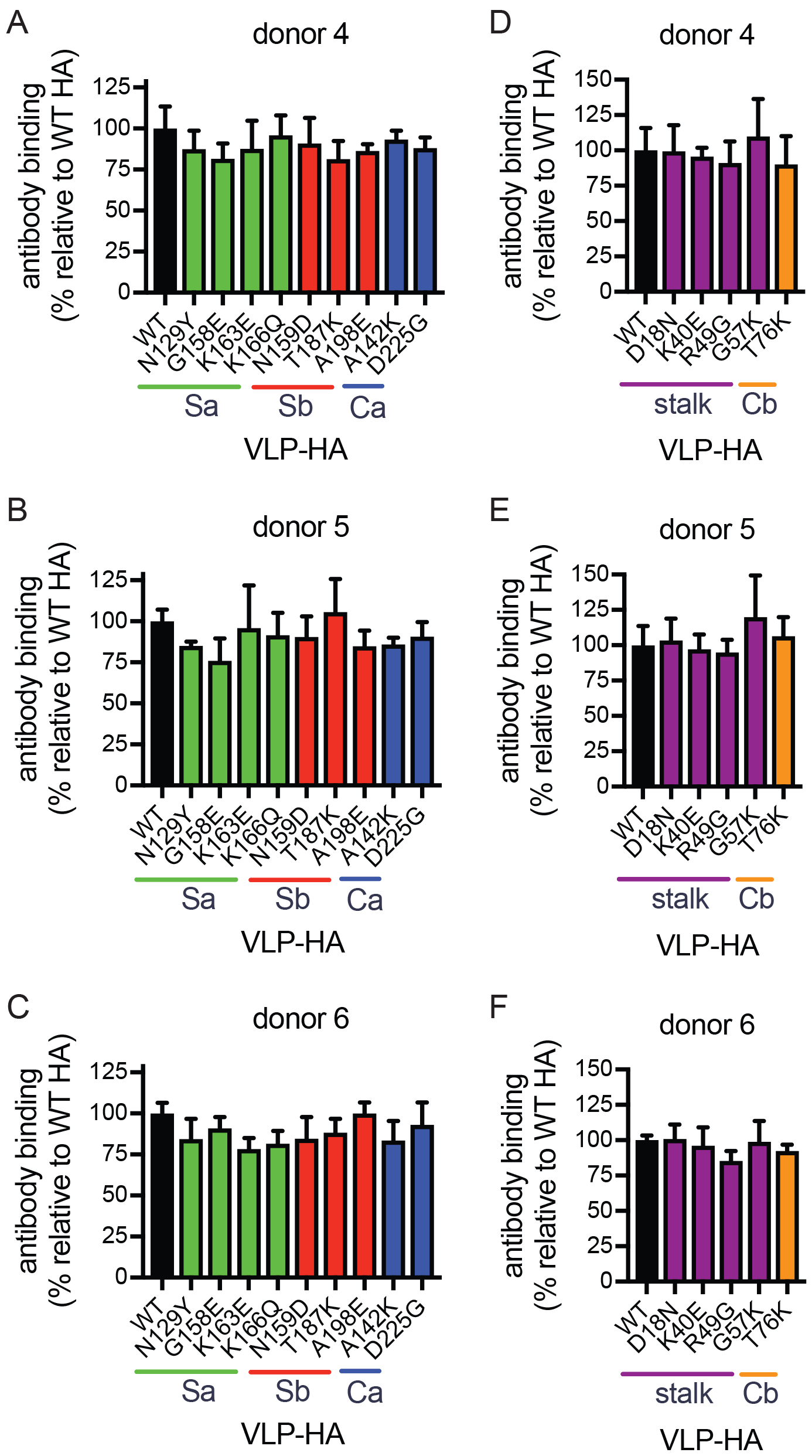
Antigenic characterization of human ‘control’ serum samples. ELISAs were completed using VLPs with HAs that had substitutions in either the top of the HA globular head (A-C) or lower part of HA (D-F). Antibody binding between different ELISA plates was normalized by including a standard monoclonal on each plate. A series of 8 serum dilutions were tested and data are expressed as % binding (based on area under the curve) relative to binding to WT HA. Experiments were completed in triplicate. Mean and SD from three different experiments are shown.

### Sequential passaging of H1N1 *in ovo* in the presence of human serum samples

To determine how H1N1 viruses evolve when confronted with narrowly focused serum polyclonal antibody responses, we passaged A/California/07/2009 virus in embryonated chicken eggs in the presence of each serum sample. As a control, we also passaged virus in the absence of serum. After the 3^rd^ passage, we extracted RNA and carried out RT-PCR and deep sequencing of each viral isolate. To reduce the influence of sequencing error in variant calling, we produced triplicate independent library preparations and averaged all replicates to determine variant frequencies. We found that viruses passaged in the presence of all three narrowly focused polyclonal serum samples acquired a substitution at >85% frequency at HA residues 166 or 167 (Fig. 5A). Residues 166 and 167 are likely in the same epitope based on their proximity (Fig. 5C), which is further supported by recent co-crystal structures of a monoclonal antibody with similar binding patterns (18). To investigate intra-sample variability, we repeated two additional viral passaging experiments with serum from donor #1. Viruses again acquired substitutions at HA residue 166, but the specific amino acid change was different in the different experiments (Fig. 5A). In contrast to passaging experiments involving narrowly focused sera, viruses grown in the presence of ‘control sera’ (i.e. sera not affected in HAI assays and ELISAs by the K166Q HA mutation) did not acquire substitutions in the HA epitope involving residue 166 (Fig 5B). Instead, viruses grown in the presence of these sera acquired substitutions at residues 128, 146, 186, 189, and 193 of HA (Fig. 5B-C). Virus grown in the absence of sera did not acquire HA mutations at >2.5% frequency. We detected several mutations in the other seven gene segments of passaged viruses, but there were no obvious mutational pattern that distinguished viruses passaged in ‘narrowly focused’ versus ‘control’ sera (Table S1).

**Fig 5.**
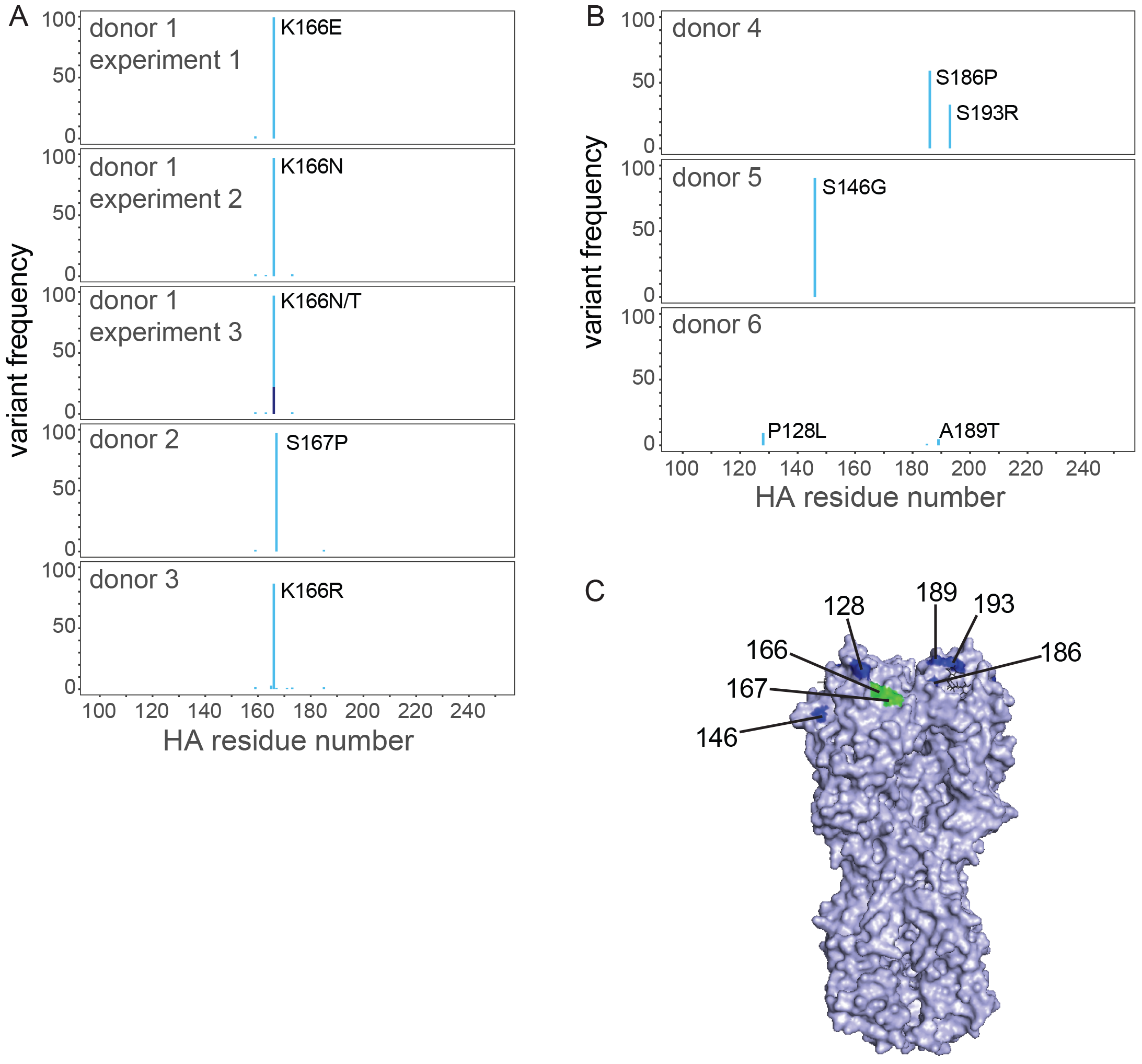
Identification of HA substitutions that arose after passaging. HA variant frequencies in virus passaged in the presence of ‘narrowly focused’ serum samples (A) and ‘control’ serum samples (B) are shown. Percentage of each variant is indicated. Different shades of blue indicate different amino acid variants at the same residue. (C) The location of HA mutations that arose after passaging are shown (PDB ID code 3UBN).

### Characterization of individual mutants that arose during *in ovo* passaging experiments

We next created VLPs possessing HA that contained substitutions that were present at >4.5% in our *in ovo* passaging experiments. We then carried out ELISAs with these VLPs and found that antibodies in serum samples that were narrowly focused on the HA epitope involving residue 166 had reduced binding to all of the variants possessing substitutions at residues 166 and 167 (Fig. 6A-C). Antibodies in these serum samples did not have reduced binding to VLPs possessing substitutions that arose in the presence of control sera (Fig. 6A-C), with the exception of VLPs possessing a substitution at residue 128, which is near residue 166 of HA (Fig. 5C). Substitutions at residues 166 and 167 also severely reduced HAI titers of these ‘narrowly focused’ serum samples (Table 1). In contrast, antibodies in our control serum samples did not have reduced binding to VLPs possessing HAs with substitutions at residues 166 and 167 (Fig. 6D-F) and did not have reduced HAI titers to viruses with these substitutions (Table 1). Antibodies in serum from donors #5 and #6 showed no reduction in binding to VLPs possessing HAs with substitutions that were selected by each autologous serum sample (Fig. 6E-F), and no reductions in HAI titers to these viruses with these mutations. Antibodies from donors #4 and #5 bound less efficiently to HAs possessing substitutions that arose during passage in the presence of donor #4 sera (Fig. 6D); however, these substitutions did not dramatically reduce HAI titers (Table 1). These data indicate that the serum samples that were narrowly focused on the HA epitope involving residue 166 efficiently selected escape mutations at residues 166 and 167 that directly prevented binding of serum polyclonal antibodies, similar to what has been previously reported for monoclonal antibodies. These data also indicate that this experimental setup has the potential to elucidate the specificity of antibodies that are present in moderate concentrations in polyclonal sera, as demonstrated by passaging experiments involving sample #4.

**Fig 6.**
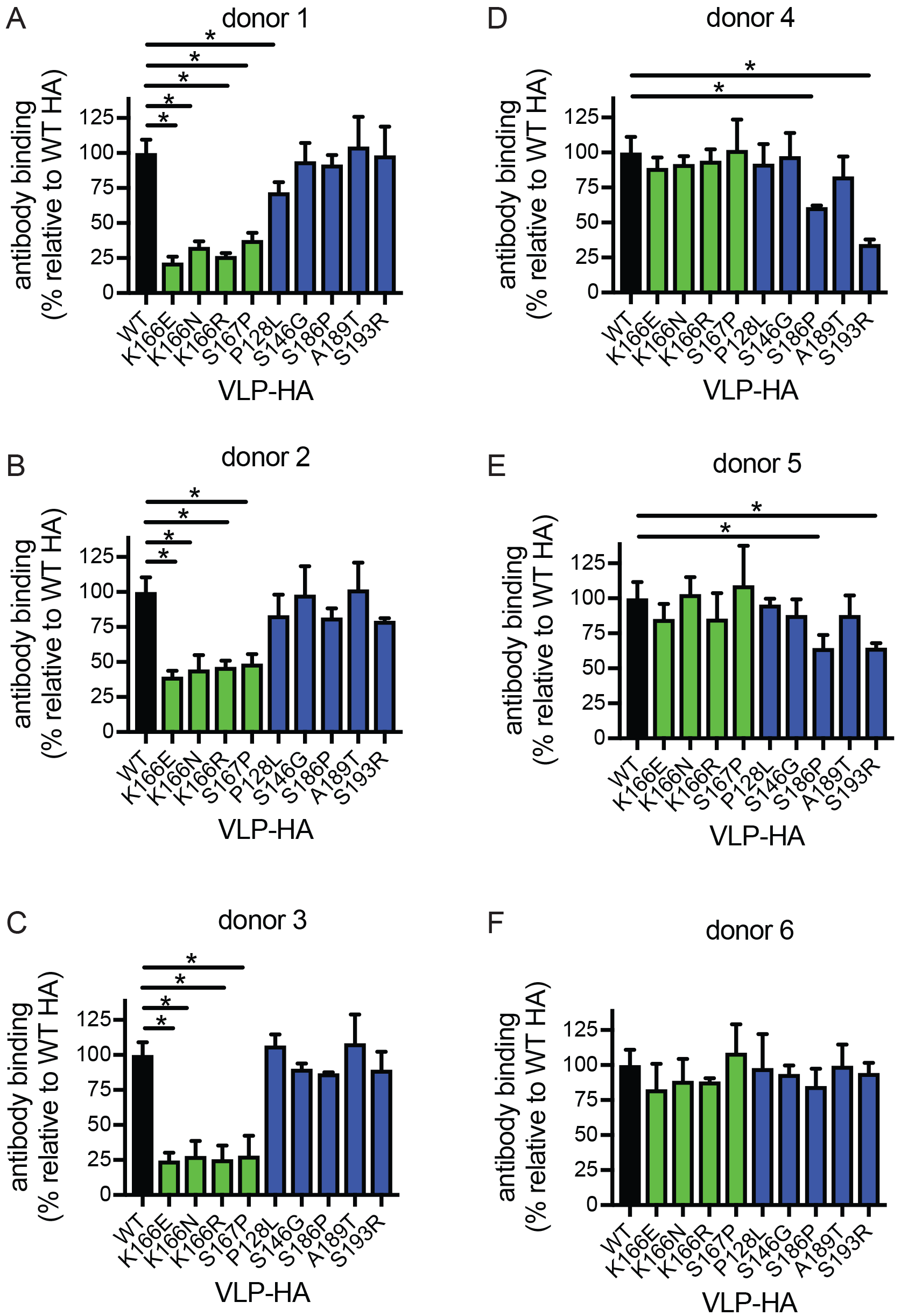
Antigenic characterization of HA variants that arose after passaging. ELISAs were completed using VLPs with HAs that had substitutions that arose after passaging in the presence of human sera. Antibody binding between different ELISA plates was normalized by including a standard monoclonal on each plate. A series of 8 serum dilutions were tested and data are expressed as % binding (based on area under the curve) relative to binding to WT HA. Experiments were completed in triplicate. Mean and SD from three different experiments are shown. (*P<0.05)

**TABLE 1.** HAI titers using viruses that emerged in passaging experiments^a^

| donor<br># | viruses |  |  |  |  |  |  |  |  |
| --- | --- | --- | --- | --- | --- | --- | --- | --- | --- |
|  | WT | K166E | K166N | K166R | S167P | P128L | S146G | S186P | A189T |
| 1 | 320 | <10 | 20 | 40 | 60 | 160 | 320 | 160 | 240 |
| 2 | 120 | 20 | 20 | 10 | <10 | 80 | 80 | 80 | 80 |
| 3 | 120 | 20 | 20 | 20 | 10 | 120 | 80 | 80 | 80 |
| 4 | 120 | 160 | 240 | 160 | 160 | 120 | 120 | 80 | 120 |
| 5 | 80 | 160 | 160 | 160 | 160 | 80 | 80 | 160 | 80 |
| 6 | 160 | 160 | 240 | 160 | 160 | 160 | 160 | 80 | 160 |

<sup>a</sup>HAI assays were completed using viruses engineered to possess the HA substitutions that arose to the highest frequency for each passage condition. Grey indicates HAI titers are decreased >4 fold relative to HAI titer using virus with WT HA.

## Discussion

Information on mechanisms by which influenza viruses evade human antibodies may be useful in understanding influenza virus evolution and forecasting future antigenic drift. In this study we studied individuals with antibody responses that were focused on a single HA epitope that was conserved in past viral strains. We found that H1N1 viruses rapidly acquired HA mutations when grown *in ovo* in the presence of these narrowly focused serum samples, and that these substitutions directly prevented the binding of serum antibodies from these individuals. Viruses grown in the presence of these serum samples acquired mutations at HA residues 166 or 167. This is notable, since these serum samples were collected prior to the 2013-2014 season, during which a mutation at HA residue 166 of H1N1 viruses quickly rose to fixation (11). A similar result using cell culture viral passaging was recently reported using convalescent sera collected in 2009 from an individual in Japan (19). We propose that influenza viruses might evolve in individuals that possess narrowly focused antibody responses by acquiring mutations that directly prevent antibody binding.

In classic studies from the 1940s and 1950s, Thomas Francis Jr. and colleagues demonstrated that influenza virus exposures boost antibody responses to viral strains that humans encounter early in life (5, 20–23). Some humans appear to have highly focused antibody responses against influenza viruses (24), and our laboratory and others have shown that H1N1 antibody responses typically recognize epitopes that are conserved between contemporary viruses and viral strains that were encountered in childhood (11, 25–27). While we have previously shown that many individuals possess H1N1 antibodies that target an HA epitope involving residue 166 (11), it is unknown if other humans possess antibodies that narrowly target other HA antigenic sites or if the majority of humans possess polyclonal responses that include many different antibody specificities. One of our ‘control’ donors (a donor that did not have HAI antibodies affected by the K166Q HA substitution) selected for viruses with substitutions at HA residues 186 and 193 that directly blocked the binding of a fraction of that donor’s antibodies (Fig. 6). This suggests that the egg passaging system described in this manuscript might be potentially useful for identifying previously undefined ‘narrowly focused’ antibody specificities.

Our study has some limitations. We completed passaging experiments in embryonated chicken eggs, but the sialic acid types that are present in eggs are much different than those in the human airway. The specific amino acid changes that arose experimentally (K166E, K166N, K166R, S167P) were different than those in variant viruses that arose in humans (K166Q, S167T). A similar observation was made by DeDiego and colleagues in H3N2 passaging experiments (28). These differences in specific amino acid substitutions could be due to differences in each variant’s ability to bind to different sialic acid types that are in eggs versus human airways. Another limitation of our study is that our mutant VLP panel was extensive but did not exhaustively cover all possible antibody binding sites on HA. Future experiments can use more recent technological advances, such as deep mutational scanning (29, 30), to better to define the specificity of serum antibodies prior to passaging experiments.

Despite these limitations, our study is important because it demonstrates that influenza viruses can escape some types of human antibodies by simply acquiring mutations that directly prevent the binding of serum antibodies. It will be important for future studies to clearly define the prevalence of ‘narrowly focused’ serum antibody responses versus more broadly reactive serum antibody responses within the human population. This information will be useful to understand which antigenic escape mechanisms (direct escape versus increased viral attachment) are preferentially used within humans. A better understanding of influenza virus antibody specificities and antiviral immune escape mechanisms will ultimately improve our ability to predict how influenza viruses might change in the future.

## Acknowledgements

This work was supported by the National Institute of Allergy and Infectious Diseases (1R01AI113047, SEH; 1R01AI108686, SEH; 1R01AI097150, ASM; CEIRS HHSN272201400005C, SEH and ASM) and Center for Disease Control (U01IP000474, ASM). Scott E. Hensley and Jesse D. Bloom hold Investigators in the Pathogenesis of Infectious Disease Awards from the Burroughs Wellcome Fund. We thank Ian York (CDC) for providing plasmids to express murine monoclonal antibodies that were used to normalize HA amounts for ELISAs.

---

**Supplemental Table 1:**
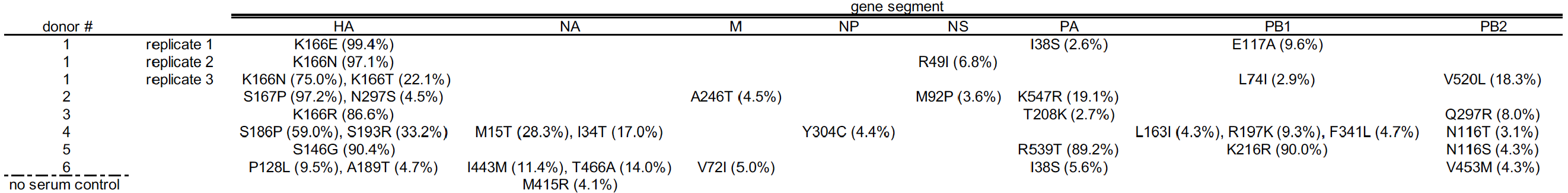
Identification of mutations in passaged viruses

